# A single-gene-based AI model to identify core and context-specific essential genes by biological interpretation from pooled genome-wide CRISPR and omics data

**DOI:** 10.1101/2024.11.03.621717

**Authors:** Chih-Yuan Chou, Jung-Yu Lee, Chia-Hwa Lee, Jinn-Moon Yang

## Abstract

Genome-wide Clustered Regularly Interspaced Short Palindromic Repeats (CRISPR) is a powerful tool for screening essential genes (EGs) and studying gene function by systematically knocking out individual genes. EGs are essential for the survival of organisms and can be divided into core EGs (CEGs) and context-specific EGs, which are crucial for the development of drugs. Although a variety of CEGs have been identified using knockout technology, the concordance among these CEG sets is extremely low; there is a lack of studies on the EG mechanisms. Therefore, developing systematic methods to identify CEGs and reveal corresponding mechanisms are important biological issues.

To address these issues, we propose a comprehensive ensembled-based model utilizing gene community-regulated pathways to decipher pan-cancer CEGs and context-specific EGs across 29 cancer types and provide insights into their regulatory abilities for each pathway. The project aims include developing a model with Systematic Identification of Essential Gene (SIEG) scores for CEGs and Context-Specific Enrichment (CSE) scores for context-specific EGs. Subsequently, we assess the regulated pathways and mechanisms of these identified EGs by integrating diverse data sources such as genome-wide CRISPR/Cas9 knockout screens, multiple omics data, KEGG pathways, and Gene Ontology. Ultimately, we aim to establish a user-friendly web service.

In our preliminary results, we gathered 1,845 genome-wide CRISPR datasets and various omics datasets, including 8,941 clinical samples and 1,346 cell lines, leading to the identification of 3,213 CEGs. By employing the SIEG score, we found that 1,178 of these pan-cancer CEGs overlapped with previously defined CEGs, reflecting a 60% similarity rate. These 3,213 CEGs play crucial roles in regulating pathways associated with cell viability, expansion, and proliferation. Additionally, they exhibit characteristics typical of CEGs, showing less favorable as therapeutic targets and centralizing within protein-protein interaction networks. Moreover, we delineated six pathway signatures of pan-cancer CEGs, encompassing transcription, translation, protein folding, replication and repair, cell growth and death, as well as energy. We anticipate that these signatures will contribute to the future redefinition of CEGs.

## Introduction

Genome-wide Clustered Regularly Interspaced Short Palindromic Repeats (CRISPR) knockout^1^ is a large-scale loss-of-function screening which provides a comprehensive view of gene function by systematically disrupting individual genes and observing resulting phenotypic changes^2,3^. This powerful gene editing technology and rapidly increasing genome-wide CRISPR knockout data offers an opportunity to systematically reveal genetic essentiality defects^2-6^. Gene essentiality refers to the fundamental role a gene plays in the viability and normal functioning of an organism^7^. Thus, a gene considered as an essential gene (EG) when its disruption significantly impairs the cell’s ability to survive, grow, or reproduce^7,8^. The identification of EGs provide crucial insights into molecular basis and biological mechanisms^9^. In general, EGs can be classified into core essential genes (CEGs) and context-specific EGs based on their essential defects across cell types or within certain context^10-13^.

CEGs constitute a group of genes deemed indispensable, the presence of which is crucial for cell viability across various tested cell lines^8,10-13^. This implies that the loss-of-function of CEGs could result in cell toxicity and undesirable side effects^12^. Consequently, recognizing CEGs is pivotal in both drug design and comprehending disease mechanisms. Besides reduced suitability as drug targets^12^, several biological characteristics of CEGs have been observed, including their high centrality in protein-protein interaction (PPI) networks^8,14-17^, high conservation, and limited occurrence of mutations^10,12^.

Compared to CEGs, context-specific EGs refer to those critical within a particular context or under specific conditions. Their context-driven specificity makes them more suitable for both diagnosis biomarkers and drug targets^10,12^. However, context-specific EGs and their mechanisms remain unclear.

To deal with the above issues, we proposed a comprehensive model by single-gene-regulated pathways from a large-scaled genome-wide CRISPR knockout corresponding with RNA sequencing data. Our model can provide a catalog of pan-cancer CEGs (29 cancer types) and cancer-type-specific EGs as well as their regulatory ability for each pathway. Based on the results, we believe that our model has shed light on core and context-specific EGs as well as their potential mechanisms.

Drawing from the listings of pan-cancer CEGs and cancer-type-specific EGs, we have confidence that our model can aid researchers in pinpointing context-specific EGs relevant to their in-house studies. By leveraging the regulated pathways provided by our model, researchers can not only acquire potential biomarkers but also uncover the underlying mechanisms at play.

## Results

### Overview

To identify pan-cancer CEGs and context-specific EGs along with their regulated pathways, we developed a systematic single gene-based pathway model. This model assigned each gene an essentiality score based on its regulated pathways (mechanisms) and a context-specificity score. Initially, we collected public genome-wide CRISPR knockout data across 29 cancer types from Project Achilles and The Biological General Repository for Interaction Datasets Open Repository of CRISPR Screens (BIOGRID-ORCS) (**Figure 1 S1-1**), and RNA sequencing data across 24 cancer types from The Cancer Genome Atlas (TCGA) and Cancer Cell Line Encyclopedia (CCLE) (**Figure 1 S1-2**). Leveraging co-expression by omics data, we quantified the regulatory strength of each gene to 342 Kyoto Encyclopedia of Genes and Genomes (KEGG) pathways called single gene-regulated pathways (GRP) method (**Figure 1 S2**). Subsequently, we utilized Systematic Identification of Essential Gene (SIEG) scoring method to prioritize each gene by their essentiality from essential and non-essential weighted pathways. We identified and evaluated a catalog of pan-cancer CEGs (**Figure 1 S3**). Leveraging gene disruption effect by knockout data, we quantified the specificity of context-dependent essentiality per gene per context called Context-Specific Enrichment (CSE) scoring method. We identified and evaluated 29 sets of context-specific EGs (**Figure 1 S4**). Afterwards, we will provide an interactive web service for researchers to query their interesting context-specific EGs and CEGs.

**Figure 1.**
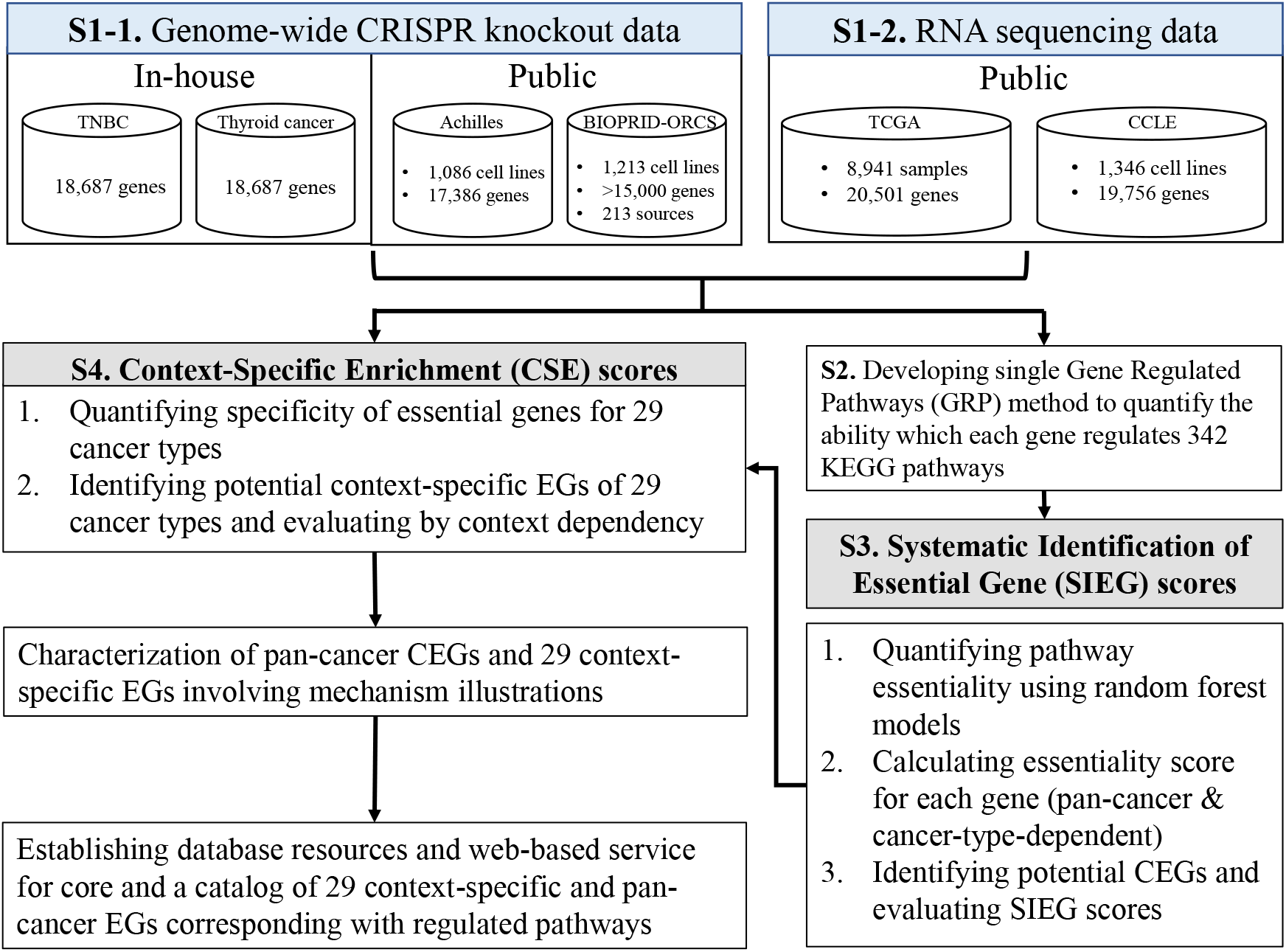
Framework of systematic single gene-regulated pathway model for pan-cancer core and context-specific essential genes. **S1**. Public genome-wide CRISPR knockout data (S1-1) is sourced from Achilles and BIOGRID-ORCS, while in-house genome-wide CRISPR data is collected from animal model-based experiments, including triple-negative breast cancer (TNBC) and highly metastatic cell lines. Public bulk RNA sequencing data sets (S1-2) are collected from TCGA (The Cancer Genome Atlas) and CCLE (Cancer Cell Line Encyclopedia). **S2**. Developing single gene-regulated pathways (GRP) method to quantify the regulated intensity per gene per pathway. **S3**. Developing SIEG scores to identify core essential gens (CEGs) and recalibrate gene essentiality by annotated CEGs regulated pathways. **S4**. Proposing CSE scores for identifying context-specific EGs to quantify the specificity of EG under certain cancer. We offer a user-friendly web service enabling researchers to acquire customized context-specific EGs based on users’ interesting.

### Identification and analysis for biological functional groups and pathway signatures of pan-cancer CEGs

SIEG scores enable the prioritization of all genes based on their essentiality, determined by0020their regulatory strength in known essential pathways. To identify pan-cancer CEGs, we selected the genes with scores greater than 1.96, totaling 3,213 genes. It indicates statistically significance in regulating known core essential pathways rather than non-essential ones. We postulated that pan-cancer CEGs contribute to regulate life maintenance pathways. Our analysis revealed a proof-of-concept that these core genes can be classified into five distinct biological functional groups based on various combinations of six pathway signatures (**Figure 2**).

**Figure 2.**
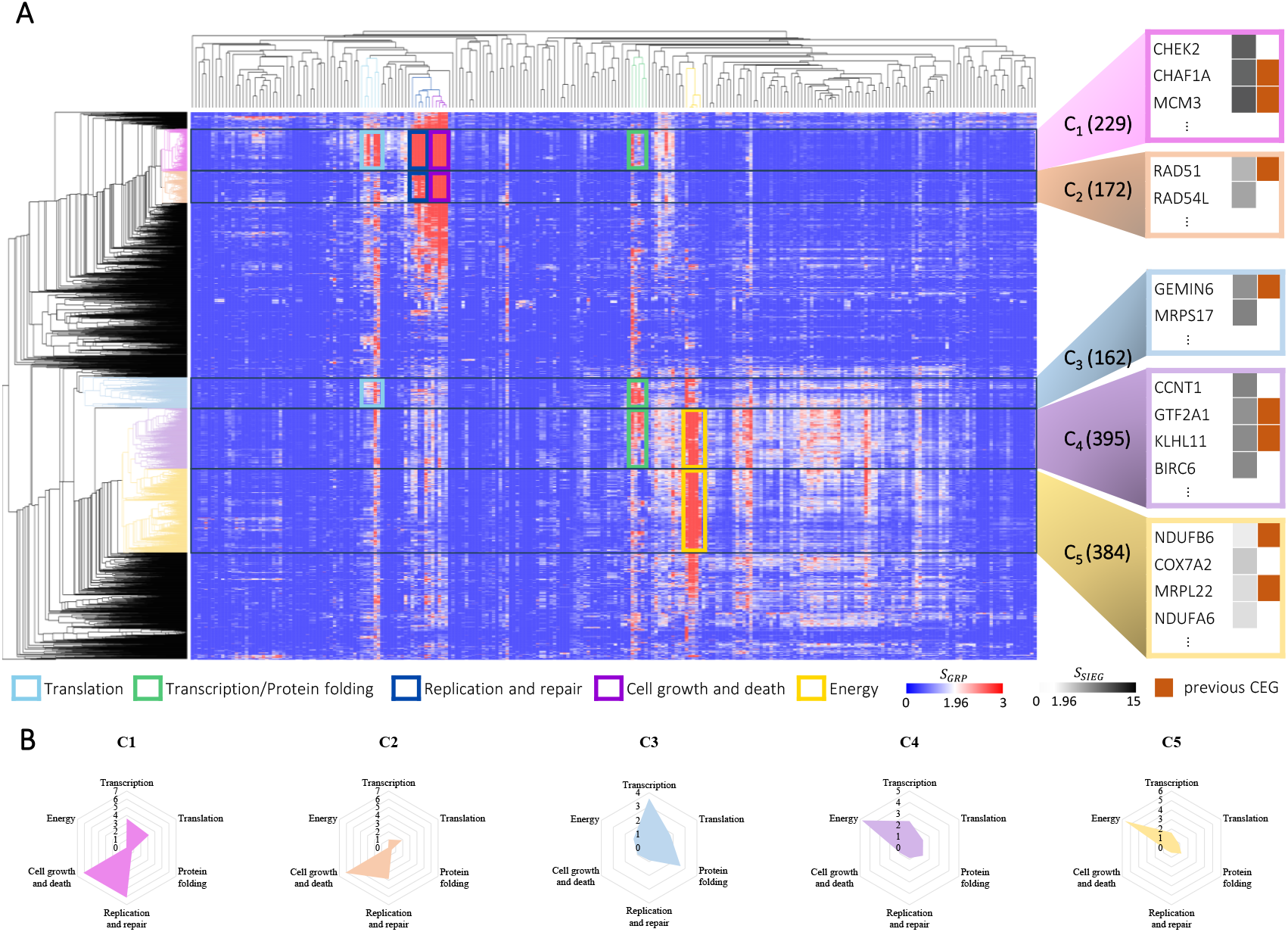
Pathway signatures of pan-cancer core essential genes (CEGs). **(A)** Six pathway signatures (biological functional groups) identified through hierarchical clustering of *z*-scores for 3,213 SIEG pan-cancer CEGs. **(B)** Gene clusters C1-C5 involve different combinations of pathway signatures. Each radar chart refers each functional group and y-axis represents averaged *S*_*GRP*_. Clusters C1-C3 regulate functions within the nucleus, while clusters C4-C5 predominately regulate energy-dependent processes. Cluster C5 includes a portion of mitochondrial genes which involve electric transport chain.

**Figure 2A** shows the hierarchical clustering analysis of 3,213 pan-cancer CEGs corresponding with 276 non-disease pathways. The pathways associated with these biological functional groups (C1-C5) exhibited a correlation with cellular viability. We further categorized each block of pathways into six pathway signatures: transcription, translation, protein folding, replication and repair, cell growth and death, and energy. The first four pathway signatures are lined to central dogma of molecular biology and occurring within cell nucleus. In contrast, the latter two pathway signatures are related to other fundamental cellular mechanisms and take place in cell plasma.

Subsequently, upon analyzing each group, we discovered that genes in group C1 regulated multiple fundamental life processes: transcription (light sage), translation (green moss), protein folding (fern green), replication and repair (evergreen), and cell growth and death (brown). C1 genes encompassed both novel and well-known pan-cancer CEGs such as CHEK2 and MCM3. Notably, MCM3 is renowned for its involvement in DNA replication and proliferation^18-20^. Through the regulation of essential pathways by MCM3, we could identify the genes such as CHEK2, exhibiting similar regulatory strength of similar pathways such as oocyte meiosis (**Figure 5B**). In contrast, C2 oversaw functions in C1 excluding transcription and protein folding. It included RAD51, whose disruption causes genomic damages, particularly in the homologous recombination repair of DNA double-strand breaks^21,22^ and cell death^22^. Based on RAD51, we then identified the genes such as RAD54L. The genes in C3 exclusively regulates central-dogma-related functions. We identified MRPS17 based on GEMIN6 which contributed to tumorigenesis^23,24^. The genes in C4 regulated not only central dogma but also energy-related functions, comprising GTF2A1 which has something to do with cell cycle^25,26^ and identifying BIRC6. In comparison with C4, the genes in C5 specifically regulated energy-related functions (**Figure 2B**), containing NDUFB6 which is involved in ROS and ATP production within mitochondria^27,28^ and identifying COX7A2.

Drawing from the regulation of diverse combinations of hallmarks, we can summarize pan-cancer CEGs and elucidate their shared and distinct essential functions, spanning cell survival, proliferation, and mitochondrial bioenergetics. Our findings underscore that SIEG scoring method can offer a detailed understanding of the underlying mechanisms of pan-cancer CEGs by utilizing expression data to explain essentiality. Hence, SIEG scoring method unveils the intricate relationship between essentiality and expression.

### Validation of SIEG scores and analysis through the biological landscape of pan-cancer CEGs

To evaluate the reliability of *S*_*SIEG*_, we validated our SIEG pan-cancer CEGs by comparing them with previously identified CEGs. Among the pan-cancer CEGs identified by SIEG scoring method, 1,178 were previously defined as CEGs, representing a 60% similarity rate (**Figure 3A**). We further assessed CEGs from the prospective of biological properties, comprising less favorable drug targets, high centrality in protein-protein interaction (PPI) networks, and a consistent tendency in consensus ratio of previous CEGs.

**Figure 3.**
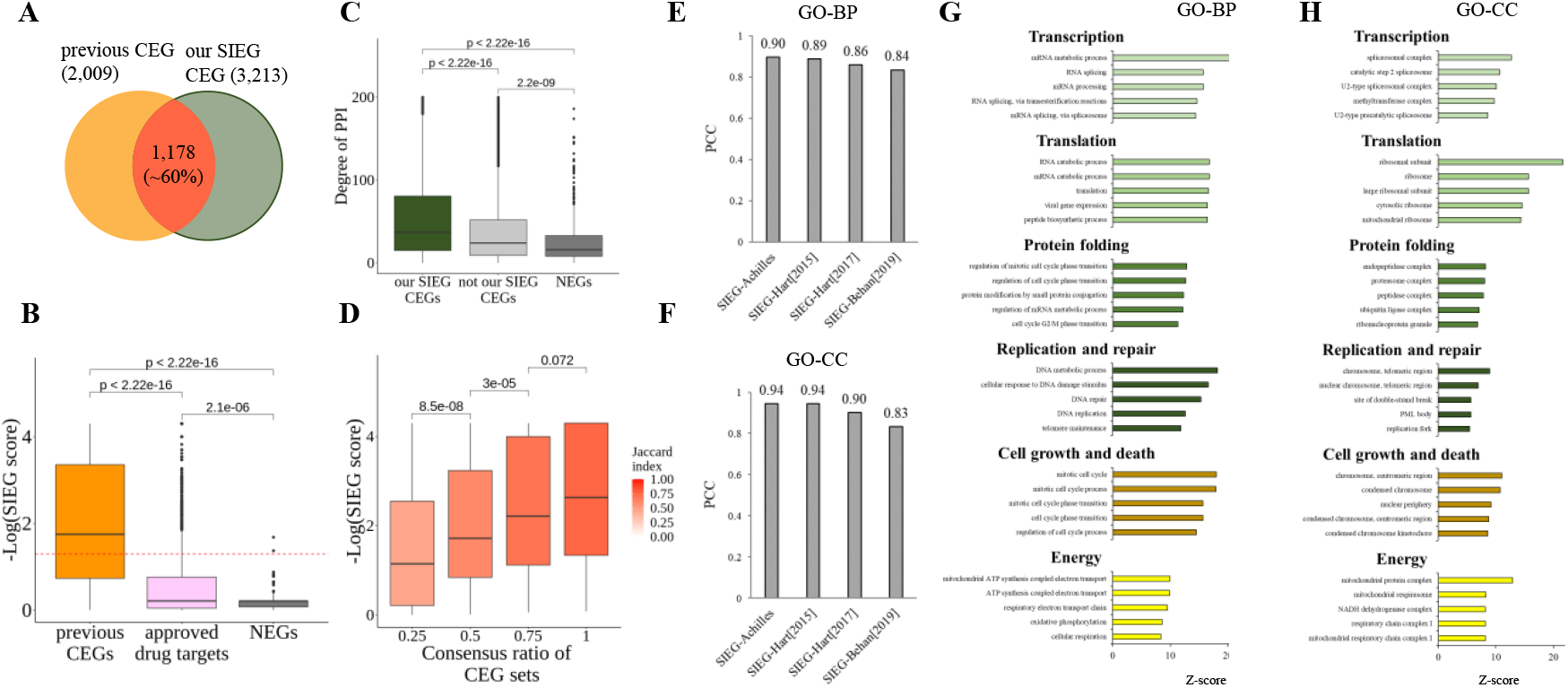
Evaluation of SIEG scoring method. **(A)** Our SIEG pan-cancer CEGs (dark green) are 60% overlapping with previous CEGs (yellow). **(B)** SIEG scores distinguish previous CEGs (yellow), FDA-approved drug drugs (pink), and NEGs (dark gray). Higher SIEG scores represent higher gene essentiality in pan-cancer. Previous CEGs are on the top. Following is FDA-approved drugs which are not suitable as CEGs. NEGs which are not essential in all cell lines are at the bottom. **(C)** Our SIEG pan-cancer CEGs show the highest degree of protein-protein interaction network. **(D)** SIEG scores are corresponding with consensus ratio of previous core EG sets. Comparing biological process **(E)** and cellular component **(F)** enrichment of our SIEG pan-cancer CEGs to previous CEGs respectively, it shows that both of them are highly similar (Pearson coefficient correlation). We further classify these biological functions and cellular components based on 6 pathway signatures. **(G)** Top 5 biological functions of each pathway signature. **(H)** Top 5 cellular components of each pathway signature.

Initially, it has been noted that cell toxicity in healthy tissues and side effects render CEGs less favorable as drug targets. However, context-specific EGs are preferred due to context-driven dependencies^12^. Consequently, we expected that previously identified CEGs would be prioritized, followed by FDA-approved drug targets, and then NEGs. Our observation indeed aligned with this anticipation. Additionally, the SIEG score distributions between previous CEGs and NEGs, CEGs and approved drug targets, as well as approved drug targets and NEGs, were found to be statistically significantly different (*p*-value determined by Wilcoxon rank-sum test) (**Figure 3B**). Secondly, it was mentioned that CEGs exhibit a greater number of interaction partners and are more likely to occupy central positions in PPI netowrks^8,10,11^. It showed that our SIEG CEGs have the highest degree of PPI and are significantly distinct from others (*p*-value determined by Wilcoxon rank-sum test) (**Figure 3C**). Lastly, our analysis demonstrated that CEGs present in more previous sets tend to exhibit higher SIEG scores and more likely to serve as pan-cancer CEGs (**Figure 3D**).

To verify the biological relevance of pan-cancer CEGs identified by SIEG scoring method, we measured the correlation of Biological Process (BP) (**Figure 3E**) and Cellular Component (CC) (**Figure 3F**) in Gene Ontology (GO) with each of four previously identified CEG sets. It was evident that our SIEG pan-cancer CEGs were highly similar with every previous CEG set in both of BP and CC, with Pearson coefficient correlations surpassing 0.8. Subsequently, we analyzed biological functions by top BPs and CCs to investigate their potential relationship to viability. We associated six hallmarks with BPs and CCs by assessing the similarity of gene members. Each BP and CC was categorized in the highest and statistically significant enrichment in a particular hallmark. Hereby, we listed top five BPs (**Figure 3G**) and CCs (**Figure 3H**) corresponding to each of hallmarks, demonstrating a high concordance with domain knowledge. Taking transcription as an example, top five BPs are related to mRNA, precisely involving the process that produces RNA from DNA. Four of top five CC terms are related to spliceosome, involving RNA splicing through removing introns from pre-mRNA transcripts. The remaining one is methyltransferase complex, involving transcriptional regulation through adding methyl groups on DNA and histone proteins.

Overall, pan-cancer CEGs identified by SIEG scores were validated through characteristics such as being less desirable drug targets, centralizing in PPI networks, and displaying a high concordance of the extent of consensus with previous CEGs. Simultaneously, they exhibited a strong alignment with the above six hallmarks in terms of BPs and CCs. This suggests that our SIEG pan-cancer CEGs outperform the others by mechanisms with biologically plausible explanations derived from multiple properties of CEGs.

### Validation of CSE scores and analysis of mechanisms and cancer dependency for cancer-specific EGs

CSE scores enable quantification of the specificity of essentiality for each gene within a given context, determined by their enrichment as EGs. Hereby, we quantified and identified 29 cancer-specific EG sets. To identify cancer-specific EGs, we considered genes with scores greater than 1.96 as cancer-specific EGs. It represents genes which are particularly essential in cell lines of the given cancer. We preliminarily identified and analyzed breast cancer, liver cancer, and thyroid cancer, which consists of 832, 483, and 42 specific EGs, respectively (**Figure 4A**).

**Figure 4.**
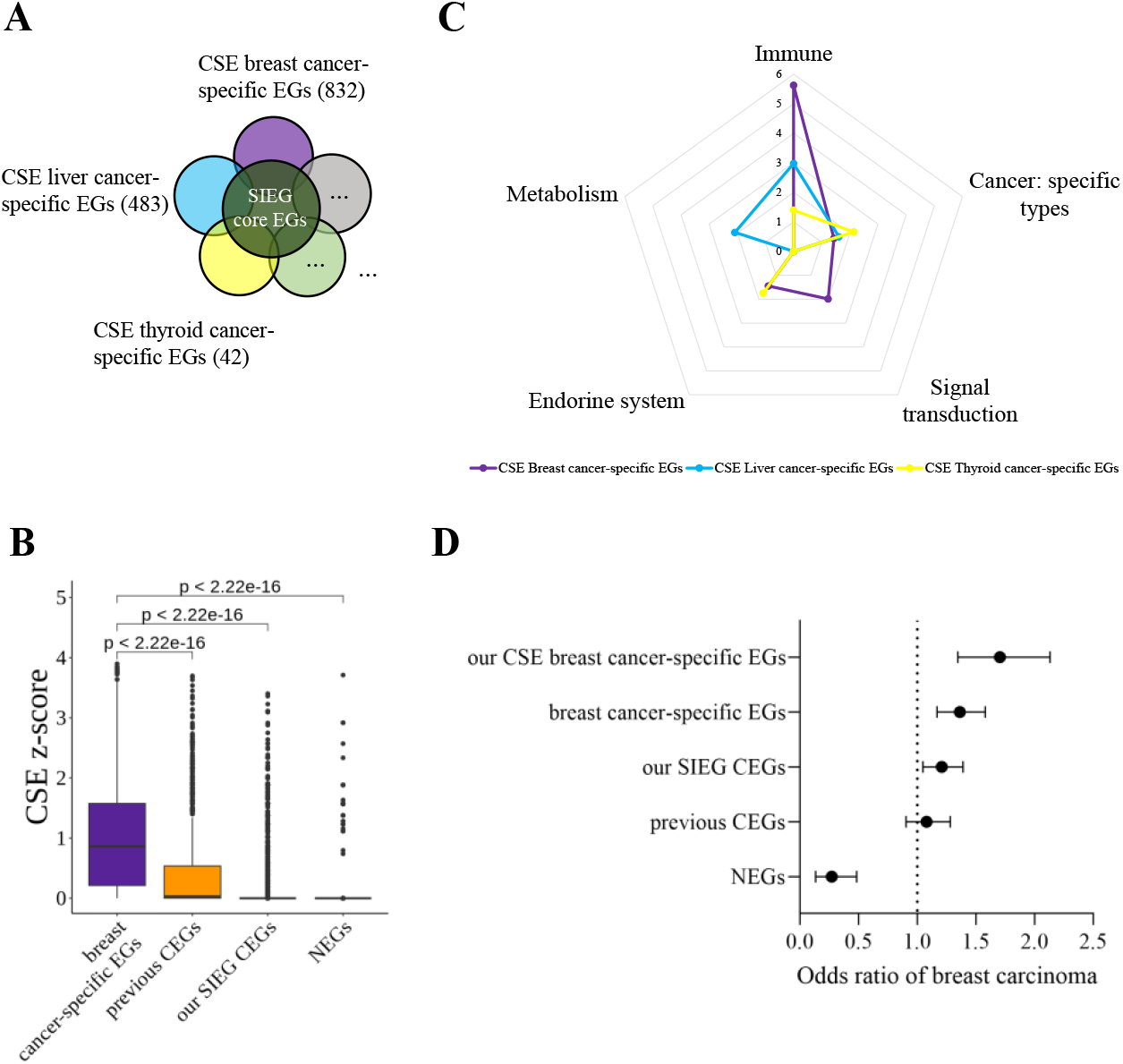
Evaluation of CSE scoring method. **(A)** Our CSE context-specific EGs, filtered by SIEG pan-cancer CEGs and other context EGs, are essential within certain context such as breast cancer-specific EGs (purple), liver cancer-specific EGs (dark blue), thyroid cancer-specific EGs (light yellow), etc. **(B)** CSE *z*-score distinguish previous breast cancer-specific EGs, CEGs, our SIEG pan-cancer CEGs, and NEGs. Higher CSE *z*-scores represent the higher specificity of EGs in a certain cancer type. In general, EGs are with higher CSE *z*-scores than NEGs. Furthermore, previous breast cancer-specific EGs are higher than both of previous CEGs and our SIEG pan-cancer CEGs. **(C)** CSE cancer-type-specific EGs display unique and common functions. Three cancer types regulate immune-related pathways; however, they are governed by different genes in each of cancer type. **(D)** Our CSE breast cancer-specific EGs show high dependency on breast carcinoma. We collect breast carcinoma biomarkers (6,776 genes with GDA score higher than 0.01) from DisGeNet. To quantify the dependency, we calculate the odds ratio of specific EGs and biomarkers in breast cancer. Our CSE breast cancer-specific EGs are at the top with the highest dependency (odds ratio is 1.7). The following includes two sets of CEGs and then previous NEGs, indicating weak or no association with breast carcinoma, respectively.

**Figure 5.**
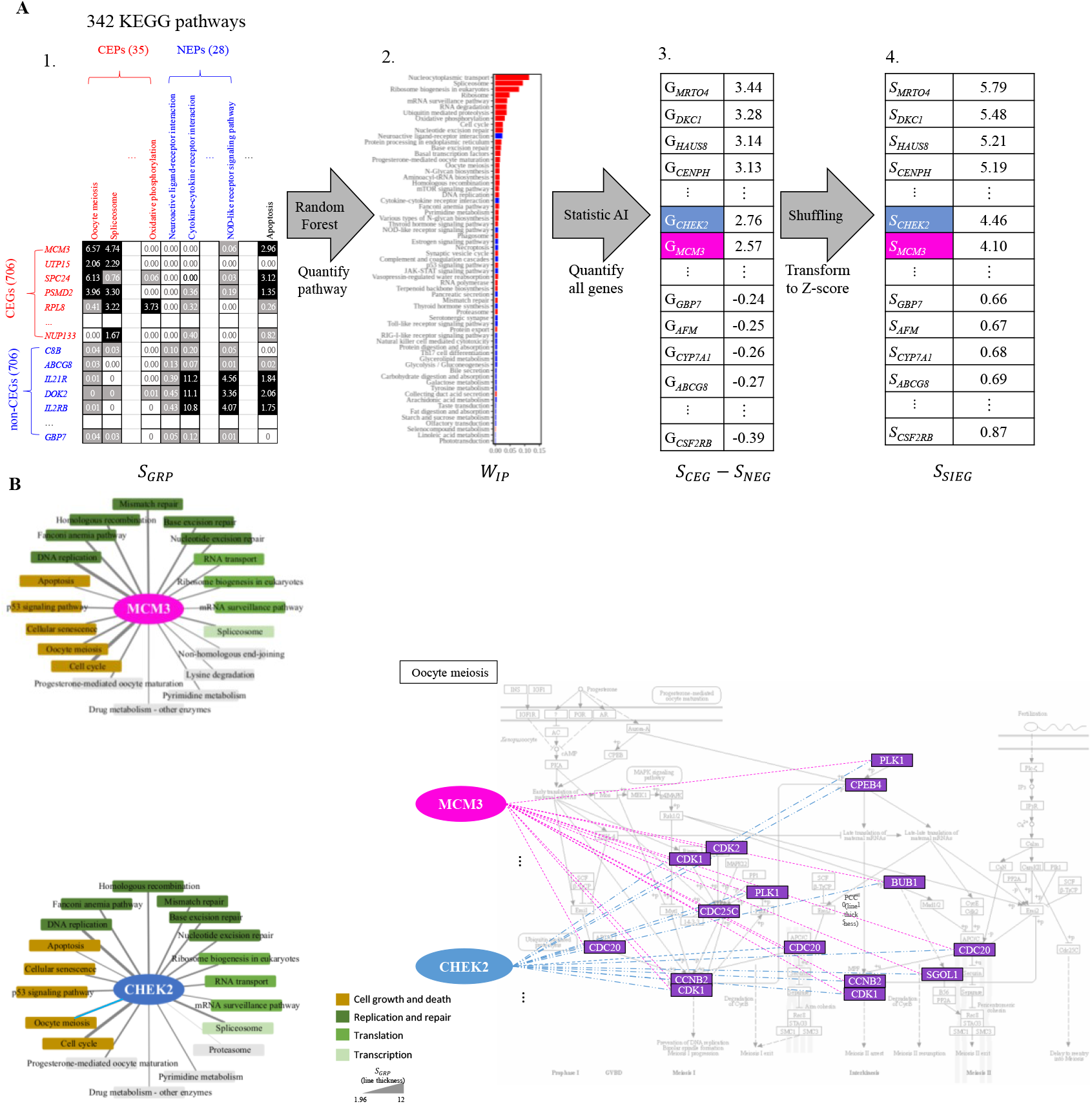
Systematic Identification for Essential Gene (SIEG) scoring method. **(A)** To quantify the regulatory capacity of each gene in modulating pathways, we calculate the enrichment of each gene’s interaction with genes in each KEGG pathway, and this measure is referred to as the *S*_*GRP*_. Quantifying pathway essentiality using random forest models is represented by the *W*_*IP*_. Calculating the regulatory capacity of gene on core essential pathways (CEPs) and non-essential pathways (NEPs), considering the statistical significance of gene essentiality through the transformation of raw scores into *z*-scores. **(B)** MCM3 and CHEK2 are identified as potential CEGs based on the *S*_*SIEG*_, with MCM3 recognized as a gene known to play a crucial role in pan-cancer. These two genes regulate similar pathways and also modulate similar genes in oocyte meiosis pathway.

To validate CSE scores, we compared our breast cancer-specific EGs to CEGs and NEGs based on their CSE scores. We hypothesized that CSE scores of our breast cancer-specific EGs would be the highest, followed by CEGs, and then NEGs. Our findings confirmed this hypothesis, and we observed that our pan-cancer CEGs identified by SIEG scores were less specific than previously identified ones. It indicated our breast cancer-specific EGs were indeed the most specific genes essential in breast cancer (**Figure 4B**).

To analyze the biological functions of context-specific EGs identified by CSE scores, we conducted the hierarchical clustering analysis for breast cancer-, liver cancer-, thyroid cancer-specific EGs corresponding with 342 KEGG pathways. We observed they had shared functions such as immune-related pathways. However, there were a distinct set of genes to regulate immune-related pathways in each cancer type. There were 25 genes (such as SLC2A3, PARP14, NFKBIE, ETS1, DOCK11, KLRG1, CASS4, CEACAM4, HAVCR2, NPL, LGALS9, CXCL11, CXCL10, and IL18) in breast cancer-specific EGs. There were 8 genes (PLCG1, SLC4A5, SLC7A8, SH2D2A, CFD, CABP4, CEBPE, and ACTB) in liver cancer-specific EGs. There were 2 genes (RAP1B and UBE2N) in thyroid cancer-specific EGs. Meanwhile, we discovered that each cancer type has their unique functions. Breast cancer-specific EGs certainly regulated specific cancer types pathways (e.g., breast cancer), endocrine-, and signaling-related pathways. Liver cancer-specific EGs regulated specific cancer types pathways (e.g., hepatocellular carcinoma) and metabolism-related pathways. Thyroid cancer-specific EGs regulated specific cancer types pathways (e.g., thyroid cancer) and signaling-related pathways (**Figure 4C**).

To further validate cancer-specific EGs based on cancer dependency, we calculated the odds ratio of context-specific EGs and disease biomarkers. An odds ratio greater than 1 represents there is connection between context-specific EGs and disease. We analyzed the relationship between breast cancer-specific EGs and breast carcinoma biomarkers. It shows that our breast cancer-specific EGs indeed have the highest odds ratio compared to others and are highly related to breast carcinoma (**Figure 4D**).

In conclusion, CSE scores revealed distinct biological functions and high context dependency. So far, we have identified 29 sets of cancer types-specific EGs and a set of pan-cancer CEGs by SIEG scores. We believe that these sets will assist researchers in identifying context-specific EGs from their genome-wide CRISPR data.

## Methods

### Systematic Identification of Essential Gene (SIEG) score development

To identify potential CEGs, we developed systematic identification for essential genes (SIEG) score (*S*_*SIEG*_) to calculate gene essentiality (**Figure 5A3, 5A4**). *S*_*SIEG*_ consists of the regulatory capacity of each gene on pathways (*S*_*GRP*_), and the essentiality of each pathway. *S*_*SIEG*_ is defined as

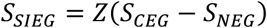

where *S*_*CEG*_ represents the regulatory capacity of CEGs on Essential Pathways (CEPs), while *S*_*NEG*_ is the regulatory capacity of NEGs on Non-Essential Pathways (NEPs). To select pathways specifically enriched in the regulation of CEGs. We subtracted *S*_*NEG*_ from *S*_*CEG*_, diminishing the significance of pathways regulated by the majority of NEGs. *S*_*CEG*_ and *S*_*NEG*_ are defined as

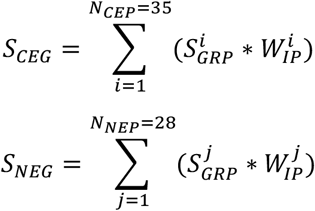

where *N*_*CEP*_ is the number of CEGs regulated pathways, while *N*_*NEP*_ is the number of NEGs regulated pathways (**Figure 5A2**). Gene Set Enrichment Analysis (GSEA) is a computational method that determines whether a pre-defined set of genes shows statistically significant and concordant differences between two biological states. We employ a previously rigorously defined set of 706 CEGs and 524 NEGs in GSEA scoring, defining CEPs when the scores of CEGs exceed those of NEGs by 1 or more, and NEPs when the scores are lower by -1 or less. According to this definition, *N*_*CEP*_ and *N*_*NEP*_ have 35 and 28 pathways, respectively, meeting the criteria. We utilize the concept of gene co-expression and the hypergeometric test to define *S*_*GRP*_, which assesses the significance of interactions between a particular gene and all the genes associated with a specific pathway (**Figure 5A1**). *S*_*GRP*_ is given as

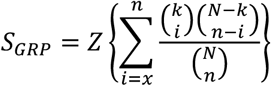

where *i* and *n* are the numbers of co-expressed gene pairs and all the combinational gene pairs in the certain pathways (e.g., oocyte meiosis), respectively; *x* is the observed co-expressed gene pairs with |Pearson’s *r*| ≥ 0.5, for instance, between MCM3 and oocyte meiosis pathway. For this example, *x* and *n* are set at 20 (indicated by pink lines) (involving genes CDC20, CDK1, PLK1) as shown **Figure 5B**, while the total number is 123. *k* and *N* are the total numbers of all the co-expressed gene pairs and combinational gene pairs, respectively, between single gene and all the genes in 342 KEGG pathways.

W_*IP*_ represents the weight of each of CEP and NEP. To assign W_*IP*_, we import Random Forest (RF) model with feature importance to quantify the contribution of each pathway in distinguishing positive and negative training sets. We use the benchmark set of 706 CEGs as the positive set. For the negative set, we randomly select 706 from a pool of 18,173 expressed genes that are not part of previous CEGs, mirroring the size of CEG training set. We input the *S*_*GRP*_ matrix, consisting of 1,412 genes and 63 pathways (*CEP*s and *NEP*s), into RF model to obtain the pathway importance of each pathway. For instance, the top three pathways include nucleocytoplasmic transport, spliceosome and ribosome genesis in eukaryotes, all intricately linked to cellular survival functions (**Figure 5A2**).

Considering the statistical significance of gene essentiality, we convert raw scores into z-scores (**Figure 5A4**). We randomly shuffle *S*_*GRP*_ matrix, which consists of 20,501 genes and 63 weighted pathways (CEPs and NEPs). We then calculate the probability of the raw score comparing to shuffling distribution. Hence, we obtain *S*_*SIEG*_, considering it statistically significant when *S*_*SIEG*_ is greater than 1.96.

Based on the analysis of *S*_*SIEG*_, we infer that MCM3 and CHEK2 could be potential CEG, with MCM3 being a known gene playing a crucial role in pan-cancer (**Figure 5B**). MCM3 was overexpressed in various human cancers, including carcinomas of the uterine cervix, colon, lung, stomach, kidney and breast^29^. Additionally, MCM3 was up-regulated in clinical human breast cancer but down-regulated after treatment^30^. Through our analysis of *S*_*GRP*_, we observed that MCM3 significantly regulates pathways such as cell cycle, DNA replication, spliceosome, and oocyte meiosis. Similarly, CHEK2 was also observed to regulate similar pathways. Furthermore, both genes similarly regulate genes (e.g., CDC20, CDK1, CCNB2, PLK1) within oocyte meiosis pathway.

### Context-specific Enrichment (CSE) score development

The wealth of genome-wide CRISPR knockout data has empowered us to systematically investigate the distribution of EGs within each context, such as cancer, a task that is not feasible with smaller-scale data. To quantify the specificity that each gene is essential in each context, we evaluated the enrichment that gene considered as EG between context and all cell lines by genome-wide CRISPR screening data across 29 cancer types from Project Achilles (1,086 cell lines) and BIOGRID-ORCS (759 cell lines). For Achilles, we deemed that genes with gene effect score less than -1 are EGs according to the median value for CEGs as -1. For BIOGRID-ORCS, which is a curated database, offers knockout profiling along with defined EGs. Here, we develop *S*_*CSE*_ score to measure context specific essentiality of each gene for a cancer type using hypergeometric distribution. *S*_*CSE*_ is defined as

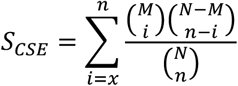

where *i* and *n* represent the numbers of essential genes in the cell lines specific to a particular cancer type and all cancer types, respectively; *x* is the observed numbers of essential genes in specific cancer cell lines; *M* and *N* refer to total numbers of specific cancer cell lines and all cell lines, respectively. Taking GSK3B in breast cancer for instance, *x* and *n* are 7 and 48; *M* and *N* are 108 and 1,827. It represents that GSK3B is essential in 48 of 1,827 tested cell lines. Meanwhile, GSK3B is essential in 7 of 48 breast cancer cell lines. Thus, *S*_*CSE*_ of GSK3B in breast cancer is 0.02. To provide essential statistics of a gene in the context specifically, *S*_*CSE*_ was transformed into *z*-score. We loosely determined the cutoff for the *S*_*CSE*_ as 1.96 for the statistical significance, aiming to obtain a set of cancer-specific EGs. Here, we acquired 29 cancer-specific EG sets for 29 cancer types. the breast cancer-specific EG set comprises 832 genes, including GSK3B.

## Notes

### Competing Interest Statement

The authors have declared no competing interest.

